# Characterising shared space use networks using animal trapping data

**DOI:** 10.1101/839530

**Authors:** Klara M. Wanelik, Damien R. Farine

## Abstract

Studying the social behaviour of small or cryptic species often relies on constructing networks from sparse point-based observations of individuals (e.g. live trapping data). Such an approach assumes that individuals that have been asynchronously detected in the same trapping location will also be more likely to have interacted. However, there is very little guidance on how much data are required for making robust co-trapping networks. In this study, we propose that co-trapping networks broadly assume that co-trapping captures shared space use (and, subsequently, likelihood of interacting), and that it may be more parsimonious to directly model shared space use. We first use empirical data to highlight that characteristics of how animals use space can help us to establish new ways to model the potential for individuals to co-occur. We then show that a method that explicitly models individuals’ home ranges and subsequent overlap in space among individuals (spatial overlap networks) requires fewer data for inferring observed networks that are correlated with the true shared space use network (relative to co-trapping networks constructed from space sharing events). Further, we show that shared space use networks based on estimating spatial overlap are also more powerful for detecting biological effects present in the true shared space use network. Finally, we discuss when it is appropriate to make inferences about social interactions from shared space use. Our study confirms the potential for using sparse trapping data from cryptic species to address a range of important questions in ecology and evolution.

**Significance statement:** Characterising animal social networks requires repeated (co-)observations of individuals. Collecting sufficient data to characterise the connections among individuals represents a major challenge when studying cryptic organisms—such as small rodents. This study draws from existing spatial mark-recapture data to inspire an approach that constructs networks by estimating space use overlap (representing the potential for interactions) from observations of individuals in the same location (e.g. a trap). We then use simulations to demonstrate that the method provides consistently higher correlations between inferred (or observed) networks and the true underlying network compared to current approaches, and requires fewer observations to reach higher correlations. We further demonstrate that these improvements translate to greater network accuracy and to more power for statistical hypothesis testing.

## Introduction

Social networks are central to addressing many of the key questions in ecology and evolution (Cantor et al. 2021). However, network construction remains a major challenge in many systems because large numbers of observations are needed to construct meaningful networks (Whitehead 2008a; Farine and Whitehead 2015). Recent technological improvements for collecting proximity, contact, or interaction data allow much more detailed networks to be constructed by improving the temporal resolution at which the data are collected (Douglas, Ji, & Clout, 2006; Rutz et al., 2012; Ryder, Horton, van den Tillaart, Morales, & Moore, 2012; Berkvens, Olivares, Mercelis, Kirkpatrick, & Weyn, 2019). However, for many smaller or more cryptic species, where observation remains difficult, many studies still rely on collecting data by trapping individuals and inferring direct social contacts from observations of different individuals occurring in the same trap at different times (i.e. shared trapping events; e.g. Perkins, Ferrari, & Hudson, 2008; Porphyre, Stevenson, Jackson, & McKenzie, 2008; Grear, Perkins, & Hudson, 2009; Grear, Luong, & Hudson, 2013; VanderWaal, Atwill, Hooper, Buckle, & McCowan, 2013; Davis et al., 2014). To date, no study has quantified whether the sparse observations typical of such studies allow us to construct robust social networks (an issue highlighted in relation to disease transmission; Tompkins, Dunn, Smith, & Telfer, 2011; White, Forester, & Craft, 2017) and, therefore, whether we can extract meaningful biological relationships from these networks.

Co-trapping networks can be constructed using a range of different data-collection methods that record shared trapping events, most commonly live-capture traps and camera traps, but also increasingly using RFID detections (Sabol et al. 2018). These methods are characterised by typically detecting single individuals at any one time, but having the capability to observe multiple individuals in the same location(s) over time. Individuals that are then observed (trapped) at the same location are considered to be connected, with binary edges (there or not) between them, or, if constructing a weighted network, with the number of shared trapping events or number of shared locations defining the strength of their connection. In co-trapping networks, two individuals that are detected in the same location are considered more likely to interact or have had direct contact (the validity of either of these assumption will depend on the biology of the system; Farine 2015). However, how much do we learn from linking (asynchronous) shared trapping events to direct contact among individuals?

The ability to construct robust networks of direct contacts will heavily depend on how well the shared trapping events at the specific trapping locations generalise to the patterns of contact between two individuals away from trapping locations. On the one hand, observing two individuals in the same trap provides some certainty that they have the possibility of co-occurring (at least within some informative timeframe). On the other hand, such observations could easily over-estimate the chance of contact between individuals. Take, for example, a trap that sits at the vertex of two individuals’ home ranges, detecting both individuals. If these are also detected once each at two additional traps in each of their respective home ranges, then we would infer an edge weight of 0.2, whereas in reality the actual area of overlap (and therefore the chances of observing them both at the same location at the same time) is negligible. Thus, co-trapping networks may be prone to error at low trapping rates (Gilbertson et al. 2021).

Constructing a co-trapping network from sparse data tacitly invokes the assumption that repeatedly observing two individuals in the same trap (at different times) informs us about their broader likelihood of coming into direct contact outside of the direct trapping observation. In other words, the resulting networks can only approximate direct contact. This raises the additional question of whether explicitly modelling shared space use might provide more powerful and robust approach to approximating direct contact. In shared space use networks, the trapping data are used to infer the amount of spatial overlap between individuals, where edge weights range from 0 if two individuals have no spatial overlap to 1 if they completely overlap. They assume that the more overlap among individuals, both at traps and away from traps, the more likely individuals are to encounter one another, resulting in direct contact (or indirect contact through asynchronous use of the same space). Several studies have used observed space sharing events to represent both shared space use and the likelihood of social interaction. VanderWaal et al. (2013) constructed a squirrel network with individual squirrels considered in contact if they were trapped in the same trap on the same day, in order to quantify the significance of shared space use, and potentially social interactions, on the transmission of a parasite. Grear et al. (2013) constructed two chipmunk networks. The first was a network where an edge between chipmunk *i* and chipmunk *j* was defined if *j* was captured in the same or nearest traps to *i* in one or two subsequent trapping sessions; a timeframe corresponding to the larval environmental development time of a nematode parasite. The authors used this network to capture potential infectious interactions. The second was a network where an edge was defined if *j* was captured in the same or nearest traps to *i* during the same trapping session. The authors used this network to capture social contacts. Further, using simulations, Gilbertson et al. (2021) demonstrated that, in telemetry studies, shared space use networks are more powerful than detections of co-occurrences when sampling is sparse (in their case, up to 72 hours between observations of individuals), but can create overly dense networks when sampling is extremely frequent. This highlights how shared space use networks and co-trapping networks are complementary approaches—both approximate direct contact rates at low sampling effort, but could be used to test different hypotheses at higher sampling efforts (e.g. asynchronous shared space use versus direct contact, respectively).

Constructing shared space use networks takes a different order of inference to co-trapping networks. First, observations (e.g. trapping data) are used to characterise individual-level space use behaviour. Only then are these space use data linked to those of other individuals. This could have several advantages. First, given that methods for estimating space use can require as few as three observations per individual (to create a polygon), it is possible that first estimating space use and then estimating space use overlap could be a relatively powerful approach when data are limited. Second, space use networks may rely less on high trapping rates, as for example trapping two individuals with overlapping ranges in interspersed traps (but never in the same trap) can still inform shared space use, but would incorrectly suggest no contact in a co-trapping network. Despite these potential benefits, to date there has been no direct quantification of the robustness of either co-trapping or shared space use networks to different sampling regimes.

It has been suggested that the data-intensive nature of networks may act as a barrier to the more widespread use of networks in the fields of ecology and evolution, with wildlife systems often being data limited (Craft and Caillaud 2011). Recent investigations into networks based on co-occurrence data (Farine and Strandburg-Peshkin 2015; Hart et al. 2021) and direct observation methods (Davis, Crofoot, & Farine, 2018) have highlighted how data-hungry networks are. Constructing a meaningful network requires sufficient observations to accurately estimate each of the many relationships (both present and absent) that connect all individuals in a population (specifically: 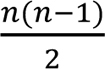 edges in an undirected network). Thus, the sampling intensity needed to maintain a minimum number of observations per individuals and, critically, the co-observations of pairs of individuals (dyads) grows quadratically with the number of individuals represented in a given network. By contrast, quantifying individual space use would result in a linear relationship between population size and sampling effort.

Previous studies quantifying the data required to construct meaningful social networks from co-occurrence data have suggested that a good rule-of-thumb is that an average of 15 opportunities to co-observe all pairs of individuals are needed (i.e. of potential associations or interactions, which could be both individuals together or just one individual in the absence of the other; Farine & Strandburg-Peshkin, 2015; Davis et al., 2018). Importantly, more observations are required for accurately defining network structure when the differences in the relationship among dyads are more uniform through a population (Whitehead 2008a, b; Hart et al. 2021), such as we expect in less social species. However, these estimates of effort may not translate well to quantifying shared space use networks, as space use is a property that can be characterised at the level of individuals.

Addressing the question of how much data need to be collected is also crucially important because many studies that have constructed shared space use networks do not report the mean number of observations per individual (Webber and Vander Wal 2019). Furthermore, the majority of those that do have been based on relatively few observations per individual. For example, VanderWaal et al. (2013) had a mean of 11 trapping events per individual, meaning that they would have a mean of 22 potential observations from which to characterise dyadic edge weights in their network (observing two individuals apart 11 times each would give the denominator of an association index—e.g. the proportion of time individuals were associated—of 22, see Hoppitt & Farine, 2018). Given the potential sparseness of observations (e.g. trapping events), then detecting two individuals at the same trap on different days is likely to capture very little information about the direct contacts between those individuals. Understanding how the number of trapping events relates to the robustness of network estimates remains a major gap in knowledge.

The choice of which data to include when constructing networks from asynchronous observations of individuals in space (e.g. trapping data) can also have an impact on how meaningful the resulting network is. Many studies have used shared space use networks to study the potential for indirect transmission events. These have typically defined a temporal threshold within which the observation of two individuals in the same place must occur for these observations to be counted as a connection (i.e. a space sharing event). The choice of threshold is directly inspired by biology, commonly the lifetime of a disease vector or pathogen in studies using networks to characterise indirect disease transmission. For example, Porphyre et al. (2008) used 28 days (maximum survival of *Mycobacterium bovis* in the environment) and Perkins, Ferrari & Hudson (2008) used 14 days (the time needed for an infective L3 larval stage of *Heligmosomoides polygyrus* to develop from the eggs of an infected host). However, such temporal definitions can be at odds with the definition and biological motivation behind applying a network approach.

For studies that rarely observe individuals, but where individuals have relatively stable ranging areas, a shared trapping event is most powerful when used to define the potential for individuals to co-occur (and possibly encounter one another) anywhere within their respective home ranges, rather than whether they actually did co-occur (and possibly encounter one another) at the particular location where the trap was set. Some studies can regularly observe or recapture individuals (such as Smith et al., 2018 who used PIT-tag readers at the entrance of burrows), and are therefore able to directly relate observed space sharing events (e.g. within a burrow on a given visit) to indirect or direct contacts. This is because observations of individuals, and subsequently space sharing events, are occurring at the same spatial and temporal scales as social behaviours or disease transmission events that form part of the study of interest (see Farine, 2018 for further discussion). Restricting the temporal scale for defining connections also reduces the data available from which the strength of connections between individuals can be estimated, which is counter-productive when animals are relatively sedentary. For example, a study that traps individuals once every few months, and discards a detection between two individuals in the same trap 15 days apart because of a maximum 14 day transmission period, would be putting too much certainty on the time-gap in detections relative to the certainty they have in terms of detecting individuals in the first place. Using observations spaced more widely apart in time forms connections between individuals that describe the system more generally as opposed to quantifying with precision the connections among specific individuals (although ideally these would correlate given sufficient observations, e.g. Sabol, Solomon, & Dantzer, 2018).

In this study, we conduct a quantitative evaluation of the robustness of co-trapping and shared space use networks to different sampling regimes. We first use empirical data to highlight that characteristics of how animals use space can help us to establish new ways to model the potential for individuals to co-occur (and potentially encounter one another). We then describe a new method for characterising shared space use network that more generally estimates home range overlap. We show that using this method can generate a network which generally (1) is more strongly correlated with the true shared space use network; (2) is a more accurate representation of the true space sharing network and, therefore; (3) has greater power to detect biological effects present in the true shared space network, relative to networks generated from the observed space sharing events only. Importantly, the spatial overlap method requires many fewer observations than using space sharing events to reconstruct meaningful shared space use networks, and many fewer observations than what has been suggested in the more general guidelines for social networks (e.g. 15 observations per dyad, see above). We also confirm that the approach is relatively robust to the underlying home range characteristics of animals. Finally, we discuss the topic of inference from shared space use networks, and how appropriate it is to link network data with biological processes.

## Materials & Methods

Our study consists of three core components. First, we use a large empirical dataset to highlight characteristics of how animals use space, specifically that they have a core and a periphery to their home range. Second, we estimate the ability for data on space sharing events by individuals to generate networks that are robust to different sampling regimes using simulated data. Third, we describe a new method, inspired by the core-peripheral nature of animal home ranges, for defining network edges. We use the same simulated data to show that this method generates observed networks which are, for a given number of captures per individual, more strongly correlated with the true shared space use network, are a more accurate representation of the true shared space use network, and are more powerful at detecting biological effects present in the true shared space use networks (i.e. detect a positive correlation present in the network from which a given simulated dataset is based) relative to networks generated from observed space sharing events only. We further demonstrate that shared space use networks can be constructed using different methods for estimating individual space use with resulting network varying in their performance, and that our findings are robust to our modelling assumptions. All simulations were run in R version 3.3.1 (R Core Team 2016) using the packages vegan version 2.4-3 (Oksanen and et al 2017) and sna version 2.4 (Butts 2016).

### Modelling home ranges in an empirical dataset

There is a large body of literature on how best to model animal space use, and it is widely accepted that many animals have a core and a periphery to their home range (Hayne 1949, 1950; Calhoun and Casby 1958; Jennrich and Turner 1969; Schoener 1981; Swihart and Slade 1989; Spencer et al. 1990; Slade and Russell 1998; Zamora and Moreno-Amich 2002; Klein and Cameron 2012). We test whether this holds true in a large-scale empirical dataset for a population of field voles (*Microtus agrestis*), and in so doing, the utility of this simple representation of home range for modelling the space use behaviour of large numbers of animals.

We use part of a dataset from a study of *M. agrestis* in Kielder Forest, UK (55°13’ N, 2°3’ W) that involved capturing individuals using live-trapping methods. Access to the study site was provided by the Forestry Commission. Full details are given in Jackson et al. (2014). The site was monitored across two years (2009-2010) by monthly trapping sessions between February and November, and contained a live-trapping grid (0.375 ha) of 150 (10 × 15) regularly spaced traps (3–5 m intervals) placed in optimal habitat. Animals were marked with passive radio frequency transponders (AVID plc, East Sussex, UK) and monitored over time, thus providing sequences of capture and recaptures. This dataset is comprised of 347 individuals and 678 trapping events.

*M. agrestis* is a polygynous species, with strictly territorial males. Home ranges of *M. agrestis* vary across different locations, habitats and across different times of the year. A review of nine studies, all conducted in later summer, but across a range of locations and habitats and using a range of different home range estimation methods, found that female home ranges varied in size from 30 to 900 m^2^ while males home ranges varied in size from 200 to 1500 m^2^ (Borowski 2003). Large males also have the largest home ranges (Borowski 2003). In our own study population, there is evidence for differences in the degree to which large males, small males and females are discouraged by distance (Davis et al., 2014). We estimate home range parameters for females and large males (mean weight ≥ 25 g) as an example, which we go on to use in our simulations (see *<u>Simulation procedure</u>*).

### Simulation procedure

In brief, our study used the following procedure (see Fig. 1):

**Fig. 1.**
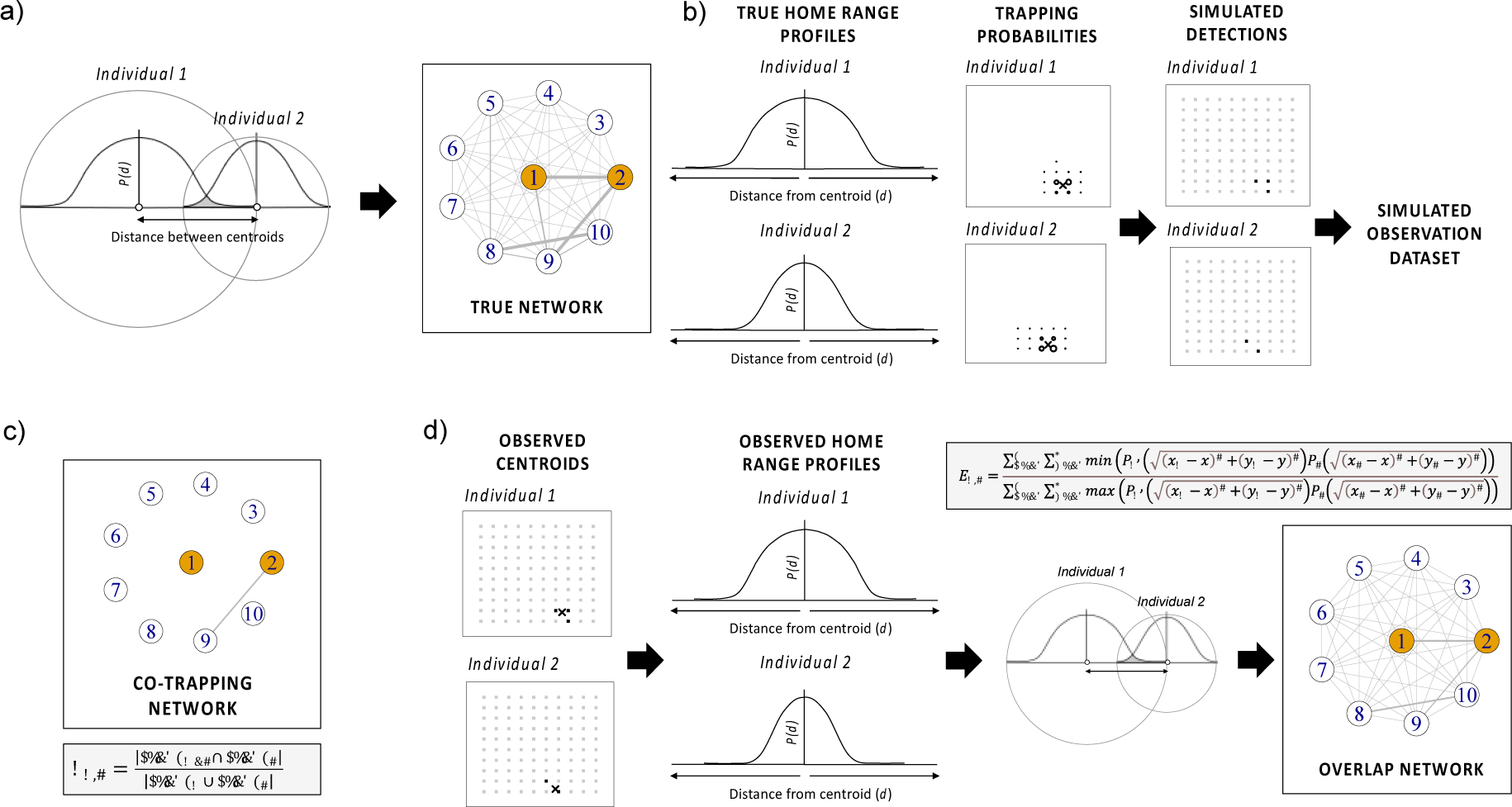
Schematic showing the simulation process: (a) Simulation of the true shared space use network with edge weights between two individuals (e.g. Individual 1 and Individual 2) equal to the overlap between their respective home ranges (here depicted along a one-dimensional slice). Each home range modelled using a negative sigmoidal curve with class-specific parameters (*a* and *b*; Equation 1) that captures the decreasing probability of observing individuals as the distance away from the centroid increases, and the overlap between the two-dimensional surfaces produced by the negative sigmoidal curves being calculated using Equation 2. (b) Generating a simulated observation dataset by calculating the probability for a given individual to be observed in a given trap based on its home range profile (crosses represent centroids; circles represent trapping probabilities; the bigger the circle, the greater the probability of detecting an individual at a trap), then simulating observations by drawing from a binomial distribution {0,1} with the probability of getting a 1 for a given individual in a given trap defined by this trapping probability. (c) Generating a co-trapping network, where nodes represent individuals, and where edge weights are calculated using Equation 3. (d) Generating an overlap network, where nodes represent individuals, and where edge weights represent the overlap between two individuals’ observed home range profiles. First, the observed centroid was calculated for each individual. Then we modelled class-specific home ranges using a negative sigmoidal curve (using a GLM regression). Third, we used Equation 2 to calculate home range overlaps as an estimate of shared space use (i.e. the edge weights), as in (a) but with the observed home range profiles and centroid values.

1. We simulated a set of 100 individuals with home ranges defined by a centroid and characterized by a negative sigmoidal curve that highlights the declining probability *P* of an individual to be detected at an increasing distance (*d*) away from the centroid of its home range:

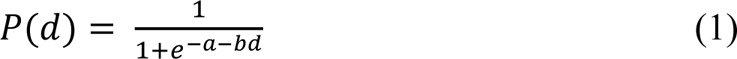

where *a* describes the overall size of the home range, *b* describes the steepness of the edge of the home range, and *d* is the logarithmic distance from the centroid. Our choice of negative sigmoidal curve was inspired by the core-peripheral nature of animal home ranges, and a large-scale empirical dataset for a population of field voles (*M. agrestis*), but we validate that our results are robust even when home ranges are defined using a uniform distribution. We defined the true shared space use network as the amount of overlap in the home range profiles across all combinations of individuals (see detailed methods below).
2. We randomly placed simulated individuals in a spatial area containing *T* traps laid out in a stratified grid. We then simulated observation datasets that contained detections of individuals at traps, where the detection probability for a given individual in a given trap was determined by the position of the trap relative to the home range profile of the individual defined in Equation 1 (higher closer to the centroid, lower further away from the centroid) or using a uniform distribution centered on the individual centroids.
3. From the simulated observation datasets, we first constructed a network based on observed space sharing events only.
4. Finally, we applied a novel method to construct an overlap network, based on estimating individual home ranges and estimating home range overlap among individuals.

Below we describe steps 1-4 in more detail:

### 1. Simulating true networks

We first drew *N* sets of *x* and *y* coordinates from a uniform distribution, where the boundaries of the distribution correspond to the edges of our study area (in our case, from 0 to 10 in each dimension). For each individual, we also randomly allocated a sex (male or female) and drew home range parameters (*a* and *b* in Equation 1) based on the sex, giving males a larger home range than females. Home range parameters for males and females were based on the empirical data (see *<u>Modelling home ranges in an empirical dataset</u>*), with added noise drawn from a normal distribution with standard deviation equal to 0.05 times the home range parameter in question (*a* or *b*) to simulate individual-level variation in home range profile.

For each simulation, we generated a true shared space use network, with edge weights representing the amount of overlap in the home range between each pair of individuals. This was done numerically by overlaying the two individuals’ 2D home range profiles and calculating the area under the two surfaces (Fig. 1a). Specifically, we predicted the probability of detecting each individual in grid overlapping both their home ranges, using equation (1), and calculated the overlap (the edge weight between individuals 1 and 2, *E*_1,2_) by dividing the sum of the lowest values at each point on the grid (*x, y*) by the sum of the largest value at each point, according to the following equation:

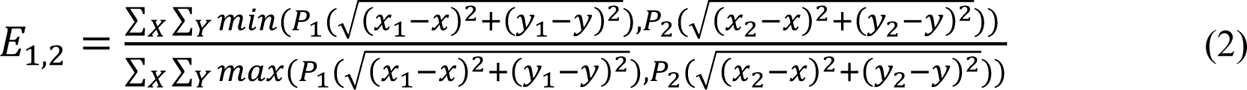

where 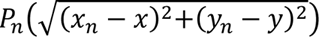 is the probability of observing individual *n*, with a home range centred at (*x*_*n*_, *y*_*n*_), at location (*x, y*) from Equation 1. X and Y represent the set of grid coordinates that encompass the home ranges of both individuals or the range of coordinates covering the entire study area, with the spacing between grid points being substantially smaller than the home ranges (e.g. every 0.1 m).

To confirm that our results are not dependent on the definition of the true networks, we also repeat our simulations by generating square home ranges with a uniform probability across their range. We do so by setting each female home range sizes to a 2.5 x 2.5 square, and male home ranges to a 3.5 x 3.5 square, roughly approximating differences in the observed empirical home ranges (results presented in Supplementary information).

### 2. Simulating observations of individuals in traps

We first calculated the probability for a given individual to be observed in a given trap. We defined this probability based on the distance of the trap to the centre of the individual’s home range using Equation 1 (or giving a uniform probability to each trap within the home range when using uniform home ranges). We repeatedly did this for all combinations of individuals and traps (‘Trapping Probability’ in Fig. 1b). We then used these probabilities to simulate observations by drawing from a binomial distribution {0,1} (‘Simulated Detections’ in Fig. 1b). We incremented the number of draws from this sampling process to generate more observations. Because draws resulted in variable numbers of observations, we then calculate the mean number of observations per individual, allowing us to make our results more easily interpretable.

### 3. Generating co-trapping networks

Each simulated dataset contained the number of detections of each individual in each trap. We generated a co-trapping network for each simulated dataset with the edge weight between individual 1 and individual 2 (*E*_1,2_) defined as:

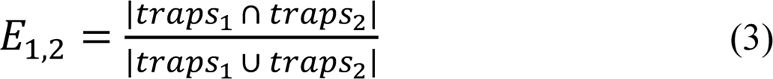

where *traps*_1_ ∩ *traps*_2_ is the set of traps in which both individuals were detected, and *traps*_1_ ∪ *traps*_2_ is the set of traps in which either or both individuals were detected (Fig. 1c).

### 4. Generating networks based on overlapping home ranges

We then applied a novel method for generating shared space use networks based on first estimating a population’s home range profile(s) from a simulated observation dataset, and then calculating the overlap in the observed home range profiles of each pair of individuals based on the distance between their observed centroids. Our method operates as follows.

First, we calculate each individual’s observed centroid by taking the mean of all of the detected locations. Second, we calculate the distance between this centroid and all of the traps where it could have been captured. Third, we calculate the observed home range profile for individuals, which (for a representative, sparse dataset) we achieve by fitting a negative sigmoidal curve (Equation 1; fitted using a Bernoulli GLM, with 0 indicating an individual was not detected at a particular trap, and 1 indicating an individual was detected at a particular trap) for males and females separately, thereby generating a relationship representing the average home range profile for each sex (see discussion for justification for this strategy of combining individuals, as well as alternative strategies). Fourth, we use the observed profiles calculated for each individual to estimate overlap in space use between each pair of individuals using Equation 2 (Fig. 1d).

We also tested whether shared space use networks are robust to different implementations by calculating home ranges using minimum convex polygons (MCPs) and estimating pairwise overlaps between individuals’ polygons (these results are presented in the Supplementary Information).

#### Estimating the robustness of shared space use networks to different sampling regimes

The ultimate aim of any network we construct is to be able to reliably test a hypothesis of interest. To test the performance of our novel spatial overlap network method against traditional co-trapping networks, we generated 1000 true networks (Fig. 1a). For each true network, we produced simulated observation datasets that varied in sampling intensity (number of draws from a binomial distribution {0,1} given a probability of observing an individual in a trap; Fig. 1b). We designed this such that the sampling intensity corresponded to a mean number of observations per individual ranging between 1 and 30 (regardless of trapping grid density), thus capturing the spectrum of what has been reported in the literature. For each simulated observation dataset, we generated a co-trapping network, and a network using the overlap approach by reconstructing separate negative sigmoidal curves (or home range profiles) for large males and females (Fig. 1c).

We assessed the performance of each of these observed networks for three metrics. First, we calculated the correlation between the edge weights in the observed network and the edge weights in the true network using a Mantel test. The correlation provides a measure of relative position of each edge, such that when the correlation is 1 the position of each edge from the observed network is the same as the positions from the true network, irrespective of any changes in scale. Second, we calculated a measure of accuracy by taking the mean of the absolute differences between the observed and true network edge weights. The accuracy provides a measure of whether the estimated edges in the observed network are on the same scale as those in the true network. Third, we calculated a measure of power by finding the proportion of observed networks in which we could detect a significant biological effect— here the difference in mean degree (sum of edge weights) between large males and females (large males were given a larger home range than females, see point 1 of the simulation procedure)—that is present in the true network (and estimated false positives by re-running the simulations without any differences between large males and females). We estimated significance for each simulated observation dataset by comparing the observed difference in mean degree between large males and females to the distribution of differences in 100 permuted networks. We used node permutations, which involved randomising the assignment of sex to the identities of each individual. We deemed the effect from an observed network to be significant if fewer than three of the randomised networks generated a difference that was larger than the observed one (two-tailed test at *p* = 0.05, see Farine, 2017).

#### Variants

We repeated the procedure described above for true networks with varying effect sizes for the difference in mean degree (sum of edge weights) between large males and females by varying the *b* parameter, resulting in: (a) an effect size half that in our empirical dataset, (b) an effect size equal to that in our empirical dataset, (c) an effect size twice that in our empirical dataset (see Results), We also repeated the procedure using trapping grids of differing densities: (a) a 10 × 10 grid, and (b) a 19 × 19 grid within the same area.

## Results

### Modelling home ranges in an empirical dataset

Consistent with space use theory, we found evidence for a declining probability of an individual field vole to use space further away from the centre of its home range.

Furthermore, we characterised this empirical relationship, between probability of detection and distance from centroid, using a negative sigmoidal curve (Equation 1; fitted using a Bernoulli GLM, with 0 indicating an individual was not detected at a particular trap, and 1 indicating an individual was detected at a particular trap). We found evidence for large males having a larger home range (*a* = 2.08, *b* = -4.82) than females (*a* = 2.83, *b* = -6.21; Fig. 2), resulting in a difference in mean degree of 1.8 between these two classes. We used these class-specific curves, and the resulting difference in mean degree, as the primary method (presented in the main text) to generate true shared space use networks in our simulations.

**Fig. 2.**
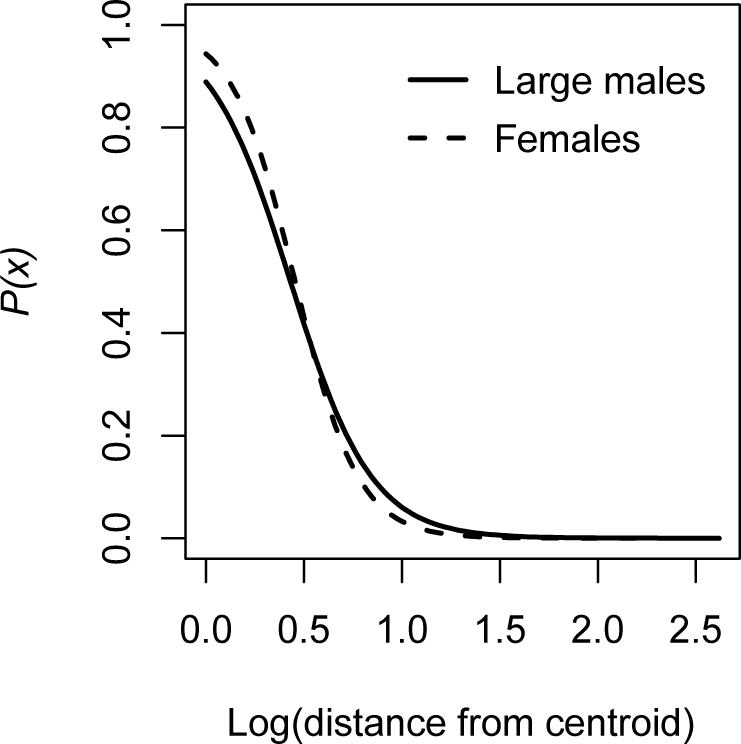
Class-specific negative sigmoidal curves for *M. agrestis* describing the change in probability of detection with increasing distance from the centre of an individual’s home range. Line shows the fitted home range profile for large males and females. Points show the raw data (whether, 1, or not, 0, an individual was detected at a location). Distances are measured in trapping grid cells (1 grid cell = 3–5 m).

### Performance of simulated observed networks with varying number of captures per individual

The number of individuals detected at least once at a trap increases with the number of captures per individual, starting from a mean of 31.3 individuals (out of a total population of 100 individuals) present at a mean of 1.3 captures per individual, and reaching a mean of 100.0 individuals (i.e. the whole population) present at approximately 10 captures per individual (Fig. 3c).

**Fig. 3.**
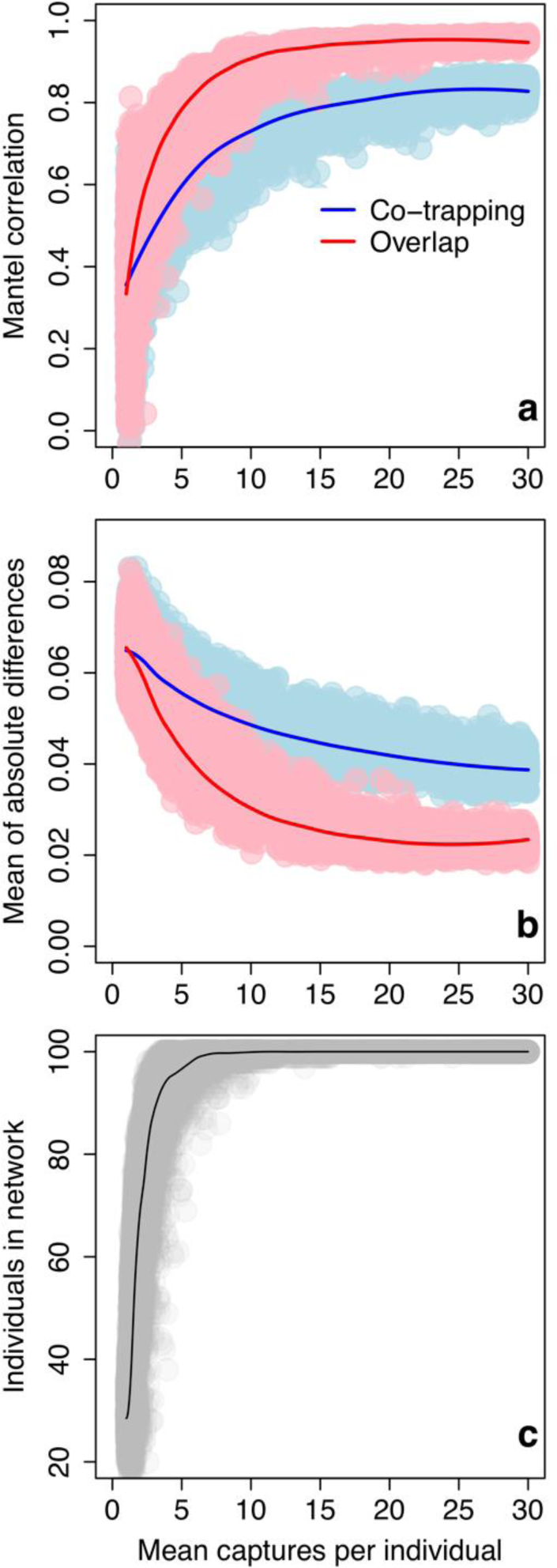
Performance of observed networks with varying number of captures per individual on a 10 × 10 trapping grid, as measured by (a) Correlation: Mantel correlation between edge weights in observed and true networks, (b) Accuracy: Mean of absolute differences in edge weights between observed and true networks (lower values = more accurate networks), and (c) Number of individuals in observed networks. LOESS smoother added to aid visual interpretation. Panel (c) refers to the data in the simulated observation dataset, which is identical for both methods.

### Correlation

As the mean captures per individual increases, the co-trapping network becomes more strongly correlated with the true network. At a mean of 1.9 captures per individual, the Mantel correlation coefficient between the co-trapping network and the true network is 0.4. The correlation coefficient plateaus from a mean of approximately 20 captures per individual, reaching a maximum of 0.8 at a mean of 28.7 captures per individual. The overlap network shows broadly the same pattern, but is, for a given number of captures per individual, typically more strongly correlated with the true network than the co-trapping network. At a mean of 1.9 captures per individual, the correlation coefficient between the overlap network and the true network is 0.5. The correlation coefficient also plateaus earlier, from a mean of approximately 10 captures per individual, and reaches a higher maximum of 1.0 (Fig. 3a).

### Accuracy

As the mean number of captures per individual increases, the co-trapping network becomes more accurate. At a mean of 1.9 captures per individual, the mean of the absolute differences in edge weights between the true space use and co-trapping networks is 6.4 × 10^-2^, which reaches a minimum of 3.9 × 10^-2^ at a mean of 28.7 captures per individual. The overlap network shows broadly the same pattern, but is more accurate for a given number of captures per individual. For example, at a mean of 1.9 captures per individual, the absolute difference in edge weights to the true network is 6.1 × 10^-2^. The mean of differences for the overlap network also reaches a lower minimum of 2.3 × 10^-2^ (Fig. 3b).

### Power

As the mean captures per individual increases, the ability to detect a true biological relationship (i.e. the power) of the co-trapping network also increases. For example, there is nearly double the chance of detecting a true positive at a mean of 4.3 captures (5.2%) compared to 1.9 captures (3.1%). However, the power remains low for small effect sizes even after large numbers of captures. The overlap network shows broadly the same pattern, but has consistently greater power to detect an effect for a given number of captures per individual, above a mean of approximately 3 captures per individual. For example, at 4.3 captures per individual there is more than double the chance of detecting a true positive in the overlap network (11.8%) compared to the co-trapping network (5.2%). The power of the co-trapping network increases continuously and reaches a maximum power of 35.2% at a mean of 28.7 captures per individual. The power of the overlap network plateaus at a mean of approximately 10 captures per individual, reaching a much higher maximum of 81.7% (Fig. 4a). Below 3 captures per individual, the co-trapping and overlap networks have similar power to detect an effect.

**Fig. 4.**
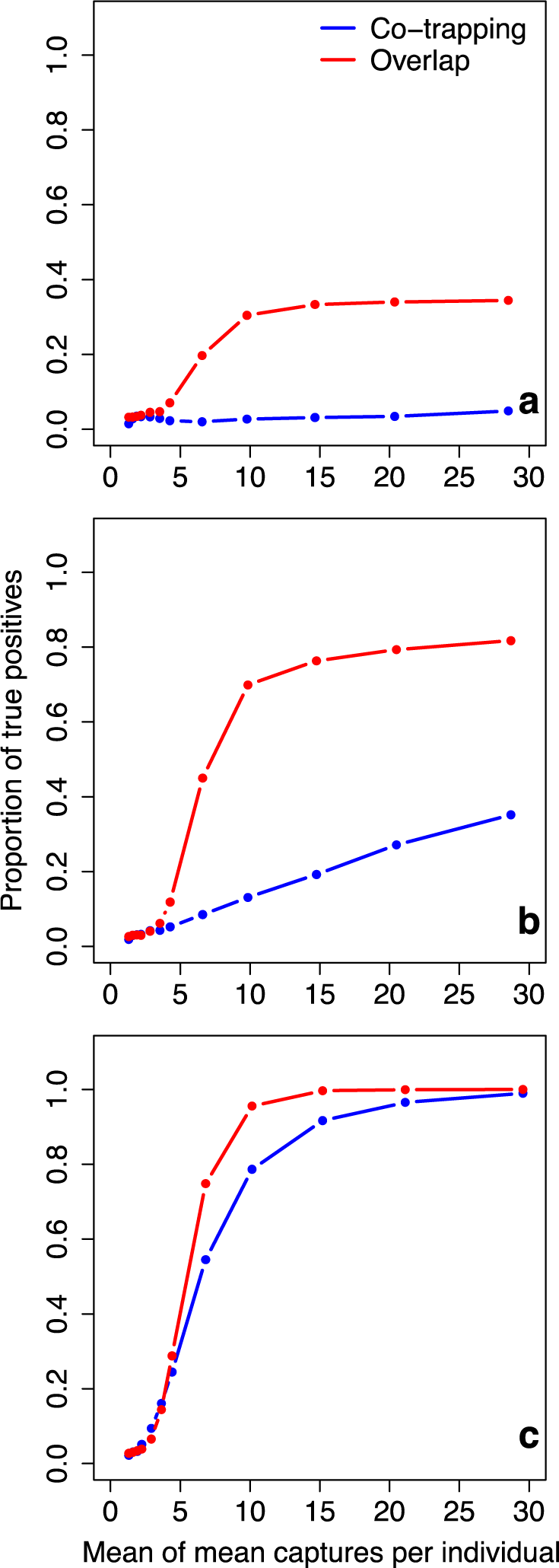
Performance of observed networks with varying number of captures per individual on a 10 × 10 trapping grid, as measured by the power of observed networks to detect a biological effect present in the true network. Proportion of true positives shown on *y*-axis, and mean of mean captures per individual shown on *x*-axis. Repeated for true networks with varying effect sizes (a) an effect size half that in our empirical dataset, (b) an effect size equal to that in our empirical dataset, (c) an effect size twice that in our empirical dataset.

Below 5 captures per individual, the co-trapping and overlap networks have a similar rate of false positives. As a mean of 9.7 captures per individual, the false positive rate of the overlap network reaches a maximum of 8%, but reduces again with more sampling. The false positive rate of the co-trapping network remains between 2% and 3%.

### Sensitivity to the modelling framework

We found that our results were generally robust to our modelling assumptions. When using a different method to simulate our true network (uniform home ranges), the overlap network remained more strongly correlated with the true network than the co-trapping network when data were sparse, and comparably strongly correlated when more data were available (see Supplementary information; Fig. S1a). The overlap network was also comparable in its power and accuracy to the co-trapping network (Fig S1b; Fig S2b; Fig S3c). However, we did not find that our results were robust to different implementations of shared space use networks. When using a different method to estimate home ranges and subsequent overlap between individual home ranges (MCPs), the overlap network generally performed less well than the co-trapping network (Fig S4) except in terms of power to detect an effect, where it performed better than the co-trapping network but less well than our own implementation (Fig S2a). The false positive rate was comparable to that of the co-trapping network (Fig S3b).

### Performance of observed networks with varying effect sizes

Only the power of the observed networks changes as a result of varying effect size.

### Power

As the effect size increases, there is a corresponding increase in the power of the co-trapping network, above a mean of approximately 3 captures per individual. For example, at a mean of 4.3 captures per individual, there is a 2.3% chance of detecting a true positive if the effect size is half that found in our empirical data, 5.2% chance is the effect size is equivalent to that found in our empirical data, and a 24.5% chance if the effect size is twice that found in our empirical data. The overlap network shows broadly the same pattern. At a mean of 4.3 captures per individuals, there is a 7.0% chance of detecting a true positive if the effect size is half that found in our empirical data, 11.8% chance if the effect size is equivalent to that found in our empirical data, and 28.8% chance if the effect size is twice that found in our empirical data. Below approximately 3 captures per individual, co-trapping and overlap networks have similar power, regardless of effect size.

### Changing trapping grid density

Observed networks change in all three metrics (correlation, accuracy and power) as a result of varying trapping grid density. Grid density also changes the number of individuals present in both observed networks when the number of captures per individual is low. For example at 1–2 captures per individual, on a 19 × 19 grid, a mean of 77.2 individuals (out of a total population of 100 individuals) are present in the observed networks (compared to 31.3 on a 10 × 10 grid; see above). However, all individuals in the population are present in the observed networks (i.e. mean of 100.0 individuals) from a mean of approximately 10 captures per individual, regardless of grid density (Fig. 5c).

**Fig. 5.**
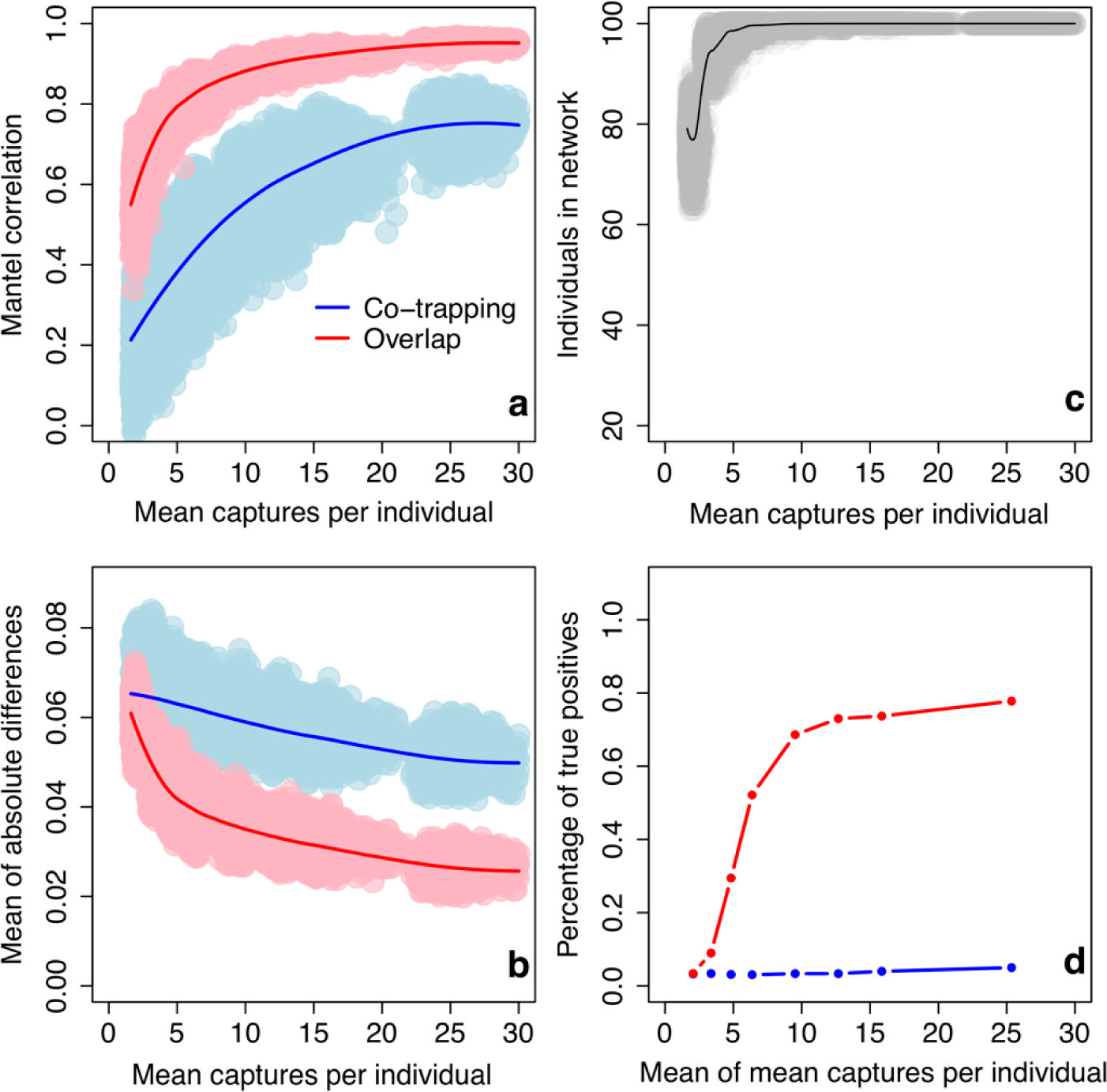
Performance of observed networks with varying numbers of captures per individual on a 19 × 19 trapping grid, as measured by (a) Correlation: Mantel correlation between edge weights in observed and true networks, (b) Accuracy: Mean of absolute difference in edge weights between observed and true networks (lower values = more accurate networks), (c) Number of individuals in observed networks, and (d) Power: Proportion of true positives. LOESS smoother added to (a–c) to aid visual interpretation. Panel (c) refers to the data in the simulated observation dataset, which is identical for both methods.

### Correlation

A higher density grid leads to a weaker correlation between the co-trapping network and the true network, for a given number of captures per individual (Fig. 5a). For example, at a mean of approximately 2 captures per individual, the Mantel correlation coefficient between the co-trapping network and the true network is 0.4 on a 10 × 10 grid, and 0.2 on a 19 × 19 grid. The co-trapping network reaches a very similar maximum correlation coefficient of 0.9 on a 19 × 19 grid and 0.8 on a 10 × 10 grid (see above). The correlation between the overlap network and true network differs very little between the 10 × 10 and 19 × 19 grid (Fig. 5a). At a mean of approximately 2 captures per individual, the correlation coefficient between the overlap network and the true network is 0.5 on a 10 × 10 grid, and 0.6 on a 19 × 19 grid. The overlap network also reaches the same maximum correlation coefficient of 1.0 on a 19 × 19 grid and on a 10 × 10 grid (see above).

### Accuracy

A higher density grid leads to a slightly less accurate co-trapping network for a given number of captures per individual (Fig. 5b). For example, at a mean of approximately 2 captures per individual, the absolute difference in edge weights to the true network is 6.4 × 10^-2^ on a 10 × 10 grid, and 6.5 × 10^-2^ on a 19 × 19 grid. The minimum absolute difference in edge weights for the co-trapping network is also higher on a 19 × 19 grid (3.3 × 10^-2^) compared to a 10 × 10 grid (3.9 × 10^-2^; see above). The overlap network shows the opposite pattern i.e. a higher density grid leads to a slightly more accurate overlap network for a given number of captures per individual (Fig. 5b). For example, at a mean of approximately 2 captures per individual, the absolute difference in edge weights of the overlap network to the true network is 6.1 × 10^-2^ on a 10 × 10 grid, and 5.8 × 10^-2^ on a 19 × 19 grid. The minimum absolute difference in edge weights for the overlap network is also lower on a 19 × 19 grid (1.9 × 10^-2^) compared to a 10 × 10 grid (2.3 × 10^-2^; see above).

### Power

The power of the co-trapping networks changes very little with grid density (Fig. 5d). For example, at a mean of approximately 2 captures per individual, the chance of detecting a true positive is 3.1% on a 10 × 10 grid, and 3.2% on a 19 × 19 grid. The power of the overlap network also changes very little with grid density (Fig. 5d). At a mean of approximately 2 captures per individual, the chance of detecting a true positive is 3.0% on a 10 × 10 grid, and 3.4% on a 19 × 19 grid.

## Discussion

In this study, we quantify the robustness of co-trapping and shared space use networks to different sampling regimes. In doing so, we provide much needed guidance for informing the choice of sampling regime when designing studies to accurately quantify space sharing among individual animals. Using a large-scale empirical dataset for a population of field voles (*M. agrestis*), we also demonstrate the utility of modelling space use behaviour on the basis that individuals have a core and a periphery to their home range. We then use these insights to develop a new method for generating shared space use networks based on estimating overlapping home ranges. We show that networks generated using the overlap method are generally more strongly correlated with the true shared space use network, are a more accurate representation of the true shared space use network, and are more powerful to detect effects present in the true shared space use network relative to networks generated from observed space sharing events only.

Our overlap method works particularly well when the mean number of captures per individual is low and provides the potential to generate meaningful networks even from sparse point-based observations of individuals. Compared to standard, more restrictive methods that rely only on joint observations at a trap and sometimes impose a temporal threshold within which the observation of two individuals in the same place must occur, our method pools data among individuals to arrive at a more general estimate of home range profile. In doing so, our method accounts for imperfect and heterogeneous observations (as in e.g. Gimenez et al., 2019). Using these general profiles, we then calculate the extent of two individuals’ home range overlap, as a function of their observed centroids, to estimate their overlap in space. Our simulation results confirm that this approach results in more accurate and more representative networks than existing methods when data are sparse, mirroring similar findings from studies simulating social contacts using telemetry data (Gilbertson et al. 2021). However, our findings are sensitive to the choice of approach. We found that overlap networks generally performed less well than co-trapping networks when MCPs were used to estimate home range overlap, except in terms of power to detect a biological effect. MCPs are a non-data-hungry method for estimating home range, but suffer from some problems. They have the opposite problem at the vertex of two individuals’ home ranges, where two individuals can be assigned an edge weight of 0 when they were trapped at the same trap and, in reality, have some non-zero level of overlap. For example, group-to-group social preferences in vulturine guineafowl (*Acryllium vulturinum*) multilevel societies are not correlated with home range overlap (Papageorgiou et al. 2019), in contrast to the relationships among superb fairy-wren groups (*Malurus cyaneus*; Camerlenghi et al. 2022). MCPs also assume a uniform density probability of occurring. We acknowledge that MCPs are overly simplistic, and encourage empiricists to find the method that best models the observed data from their animals when estimating their space use.

In our own implementation, we model differences in home range profile between large males and females based on our empirical data. However, classes of individuals which differ in their space use will vary between systems, and prior knowledge will be necessary to identify these classes e.g. males and females, larger and smaller individuals or younger and older individuals (Wolton and Flowerdew 1985; Mikesic and Drickamer 1992; Dahle and Swenson 2003; Godsall et al. 2014). Given adequate classification and modelling, our results show that an increase in correlation and accuracy of the overlap method using our own implementation translates to greater power at extracting biological effects present in the true shared space use network. In our case, we modelled differences in mean degree (sum of edge weights) between large males and females, but these outcomes should be generalisable to other hypotheses. As noted above, the overlap network generated from MCPs was also more powerful than the co-trapping network, but less powerful than our own implementation.

An important, and perhaps unexpected, finding is that denser grids of data-collection traps can make co-trapping networks based on space sharing events less strongly correlated with the true network and less accurate (at least when the mean number of captures per individuals is below 30). This result makes sense when considering sampling stochasticity. If there are more available traps, then it is less likely that two individuals, which overlap in space, will be trapped in *exactly* the same trap (unless the number of captures is very high and in no way limiting). Subsequently, constructing networks from occurrences at the same trap reduces the numerator of the edge weight calculation in Equation 3 (the set of traps in which both individuals were detected) and increases the denominator (the set of traps in which at least one individual was detected). By contrast, we show that, in terms of correlation, accuracy and power, our overlap method performs equally well, or better, on a denser trapping grid. This is because finer-scaled grids provide better estimations of individuals’ space use.

It is important to note that networks generated using the overlap method do not always perform better than those based on observed space sharing events only. We show here that a co-trapping network is more accurate and more powerful at detecting biological effects present in the true network when the effect size is low and the number of captures per individual is high (and the trap density is low). In other words, if many space sharing events are observed, then a network based on these alone deviates less from the true network and is more likely to detect a subtle biological effect present in the true network, relative to the process of pooling data and generating population-wide home range profiles. This finding aligns with the simulation results of Gilbertson et al. (2021), who found that—for telemetry data—shared space use networks also became less sensitive at higher sampling rates (albeit, much higher sampling rates than could ever be achieved from trapping data). Thus, studies that use methods that produce substantially larger datasets than singular trapping does, such as RFID detections (e.g. Sabol et al., 2018), should model the sampling process to determine the most powerful approach for a given effect strength.

We further note that the performance of co-trapping networks may also depend on the biological system, or the ecological conditions it experiences (as in Perkins, Cagnacci, Stradiotto, Arnoldi, & Hudson, 2009). Our study is based on empirical data on *M. agrestis* sampled during the breeding season. During this time, field voles maintain relatively fixed home ranges which can be estimated with some certainty (Myllymaki 1977; Niethammer and Krapp 1982). However, this is likely to be problematic if individuals are highly mobile, resulting in constantly shifting home ranges. We would expect a co-trapping network to be better suited to more dynamic systems. Our study population also inhabits a relatively homogenous landscape, in the form of grassy clear-cuts within a coniferous forest. As a result, individuals are not expected to vary a great deal in the size of their home range.

Landscape features, such as hills, in a more heterogeneous landscape could result in more variability in home range size among individuals, making it difficult to quantify an ‘average’ home range. If sampling is sufficiently high (Noonan et al. 2019), individual differences in home range profile could be accounted for when using the overlap method. This could be done, for example, by fitting a random effect for individual within class-specific regressions. These individual home range profiles could vary in size (by changing the *a* and *b* parameters of the negative sigmoidal curve) and/or in shape (by replacing the negative sigmoidal curve with a different function or an explicit home range model, e.g. Fleming & Calabrese, 2017). A number of approaches, beyond MCPs, also exist for modelling home ranges that could be employed (Winner et al. 2018). However, a co-trapping network could be better suited if aiming to study direct social contacts given the potential loss of performance from shared space use networks at high sampling frequencies (Gilbertson et al. 2021)

Shared space use networks are, and will continue to be, widely used to shed light on various biological processes. For example, individuals who share more space may be more likely to compete for resources. Many parasites and pathogens are also transmitted through the environment, and so knowing who shares space with whom can tell us something about who is likely to transmit infection to whom (VanderWaal et al. 2013). It is also true that shared space use, or proximity, is a prerequisite for interaction (Farine 2015), but whether or not individuals that share space do indeed associate, or interact, and thus how far point-based observations can be used to draw meaningful inferences, will depend on the biology of the system. Behaviour in particular is important to consider, with some animals actively avoiding each other (Davis et al., 2014) and others actively seeking each other out (Raulo et al. 2021). It is therefore important to take care when making biological inferences from any network data.

One point we highlight in our study is that the process of network generation makes explicit assumptions about the biological processes being modelled. Calculating space sharing events, for example the presence of two individuals in the same location within a given pathogen transmission period (defined by the lifetime of the pathogen in the environment, or the time taken for a pathogen to develop into an infective stage) produces networks aimed at estimating potential transmission events that actually took place (i.e. contact, which could or could not have resulted in transmission). When observation data are sparse, these observed events are likely to represent only a small proportion of all events that took place, and thus the power of the network to detect biological effects is low. This could explain why observed transmission networks are not always robust estimates of transmission processes (Wohlfiel et al. 2013). By contrast, modelling the overlapping space use among individual captures the relative probability of indirect contact (necessary for transmission events) taking place among all the dyads in a population, which will include both observed and unobserved events. Our simulations confirm that defining networks in this way can produce networks that are more powerful at detecting biological effects, especially when observations are sparse (see also Gilbertson et al. 2021). We use pathogen transmission as an example to illustrate our point, but this should be generalisable to other questions.

Our method provides a novel opportunity to generate meaningful shared space use networks, and if appropriate, to make inferences from shared space use about social interactions, even from sparse point-based observations of individuals. It therefore unlocks the potential of these data, still the most common form of data available for many smaller or more cryptic species, to address a range of key questions in ecology and evolution.

## Glossary

True shared space use network: A network which is a true representation of space-sharing amongst individuals in a population, against which observed shared space use networks are compared. Edge weights represent the amount of overlap in the home range between each pair of individuals.
Sampling intensity: The number of draws from a binomial distribution {0,1} given a probability of observing an individual in a trap. Combined with trapping grid density, results in a mean number of observations per individual.
Sampling regime: A combination of (i) the trapping grid density used in a study, and (ii) the sampling intensity in this grid, resulting in some mean number of observations per individual for a study.
Individual centroid: The centre of an individual’s home range. Calculated by taking the mean of all the positions it was observed in.
Home range profile: The change in probability of detection with increasing distance from the centroid of an individual (here described using a negative sigmoidal curve, but see Discussion). Assumes an individual has a core and periphery to their home range, with the probability of detection higher closer to the core, and lower further away from the core.
Simulated observation dataset: A dataset that contains detections of individuals at traps, where the detection probability for a given individual in a given trap is determined by the position of the trap relative to the individual’s true home range profile.
Space sharing event: The observation of two individuals in the same spatial locations, within a given defined temporal boundary (if applicable).
Co-trapping network: A network based on observed trap-sharing events only. Edge weights represent the number of traps in which both individuals were detected divided by the number of traps in which either or both individuals were detected.
Spatial overlap network: A network based on the amount of spatial overlap in the home range among individuals. Edge weights represent the overlap between two individuals’ home ranges. In our case, we calculated home range overlap numerically by overlaying the two individuals’ home range profiles and calculating the area under the two curves using Equation 2.
Observed network: A network in which the estimation of the relationships (here space-sharing) among individuals in a population is based on an observed dataset. In our case, we simulated observed datasets and created observation networks using both space sharing (co-trapping) and spatial overlap approaches.
Power of observed networks: The proportion of observed networks in which one can detect a biological effect known to be present in the true network.
Correlation of observed network: The correlation between the edge weights in an observed network and the edge weights in a true network using a Mantel test.
Accuracy of observed network: The mean of the absolute differences between the edge weights in an observed network and the edge weights in a true network.

## Acknowledgements

KMW was supported by a Natural Environment Research Council (NERC) research grant NE/L013452/1 awarded to Mike Begon, Steve Paterson (University of Liverpool), Janette Bradley (Univeristy of Nottingham) and Joseph Jackson (University of Salford), and a Johnston Postdoctoral Development Award from the University of Liverpool. DRF received funding from the Max Planck Society, an Eccellenza Professorship Grant of the Swiss National Science Foundation (Grant Number PCEFP3_187058), the European Research Council (ERC) under the European Union’s Horizon 2020 research and innovation programme (grant agreement No. 850859). The empirical data used in this study were collected as part of the NERC research grant NE/E015131/1 awarded to Mike Begon, Steve Parterson, Janette Bradley and Richard Birtles (University of Salford). We would like to thank the many individuals involved in generating this dataset, particularly Joseph Jackson, and the Forestry Commission for access to the study sites.

## Declarations

### Data availability

Data available from the Dryad Digital Repository: https://datadryad.org/stash/dataset/doi:10.5061/dryad.bk537. The R code demonstrating how to apply our method and to replicate the simulations is provided with our submission, and will be deposited in Dryad on acceptance.

### Competing interests

The authors declare no competing interests.

### Author contribution statement

Both authors designed the study, analysed the data and wrote the manuscript.

## Notes

https://github.com/kwanelik/Shared-space-use-networks

### Competing Interest Statement

The authors have declared no competing interest.

### Summary of Updates

Added testing of robustness of our results to (1) different implementations of shared space use networks, and (2) different definitions of the true networks. Summarised in the main text and in a new Supplementary information file.

