## Supplementary information for "Characterising shared space use networks using animal trapping data"

*Klara M. Wanelik*

Current address: Department of Zoology, University of Oxford, Oxford, UK.

*Damien R. Farine*

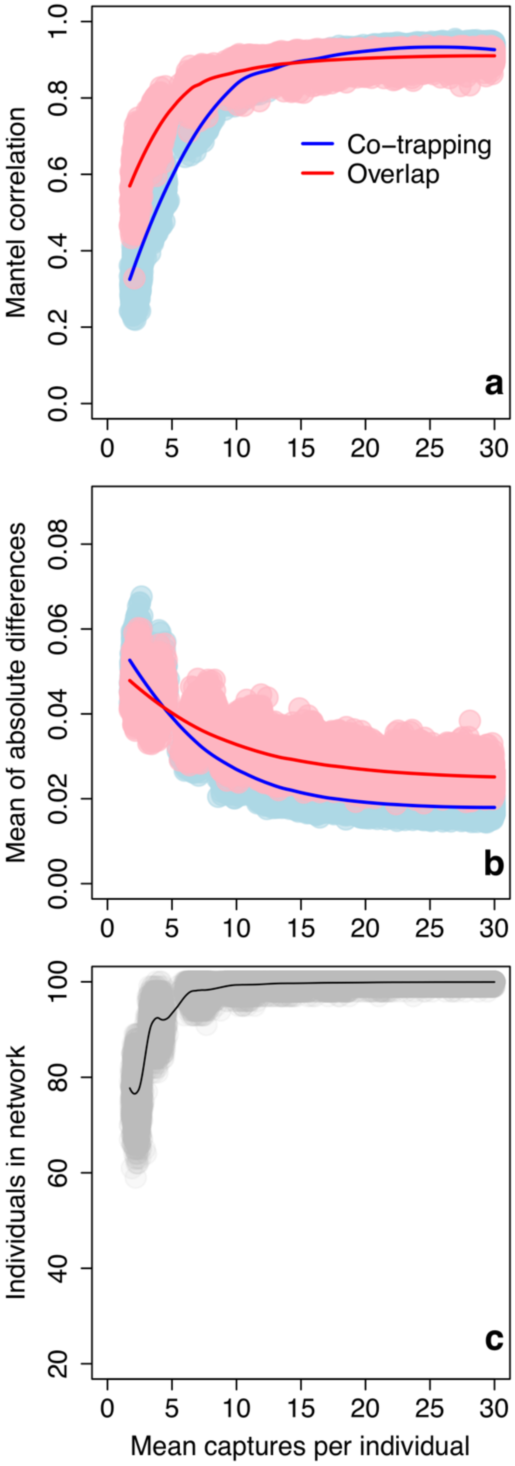


**Fig. S1** Performance of observed networks generated using our own implementation with varying number of captures per individual on a 10 × 10 trapping grid, as measured by (a) Correlation: Mantel correlation between edge weights in observed and true networks generated using uniform home ranges, (b) Accuracy: Mean of absolute differences in edge weights between observed and true networks generated using uniform home ranges (lower values = more accurate networks), and (c) Number of individuals in observed networks. LOESS smoother added to aid visual interpretation. Panel (c) refers to the data in the simulated observation dataset, which is identical for both methods.

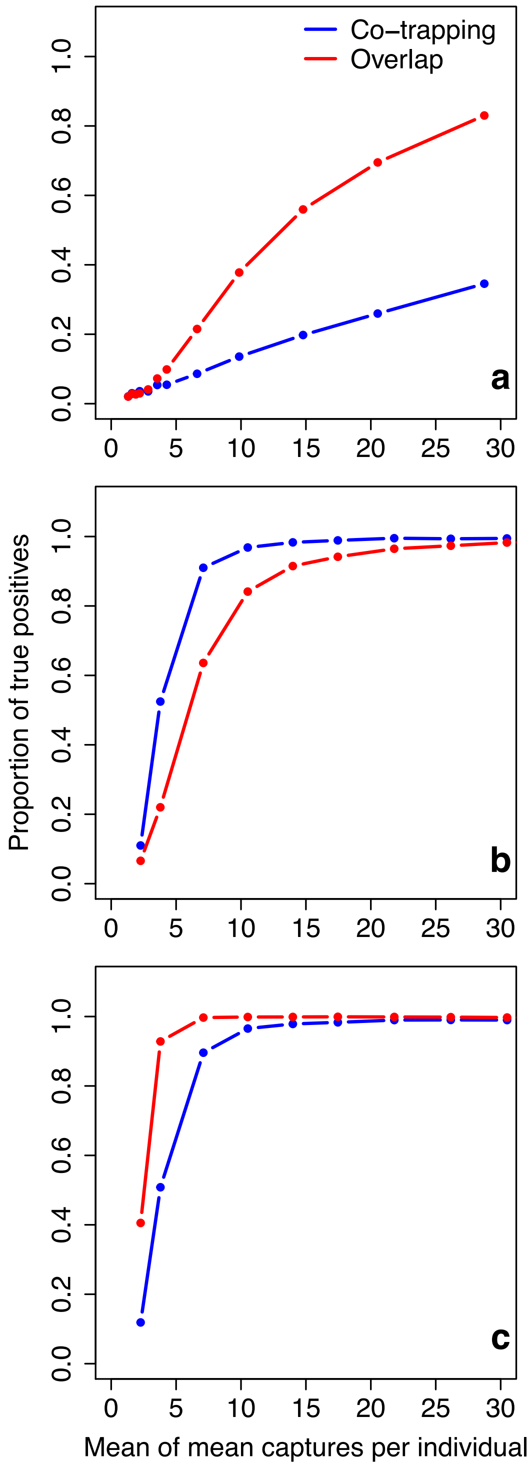


**Fig. S2** Performance of observed networks with varying number of captures per individual on a 10 × 10 trapping grid, as measured by the power of observed networks to detect a biological effect present in the true network. Proportion of true positives shown on *y*-axis, and mean of mean captures per individual shown on *x*-axis. Repeated for (a) observed networks generated using minimum convex polygons (MCPs) and true networks generated using our own implementation, (b) observed networks generated using our own implementation and true networks generated using uniform home ranges, (c) observed networks generated using MCPs and true networks generated using uniform home ranges.

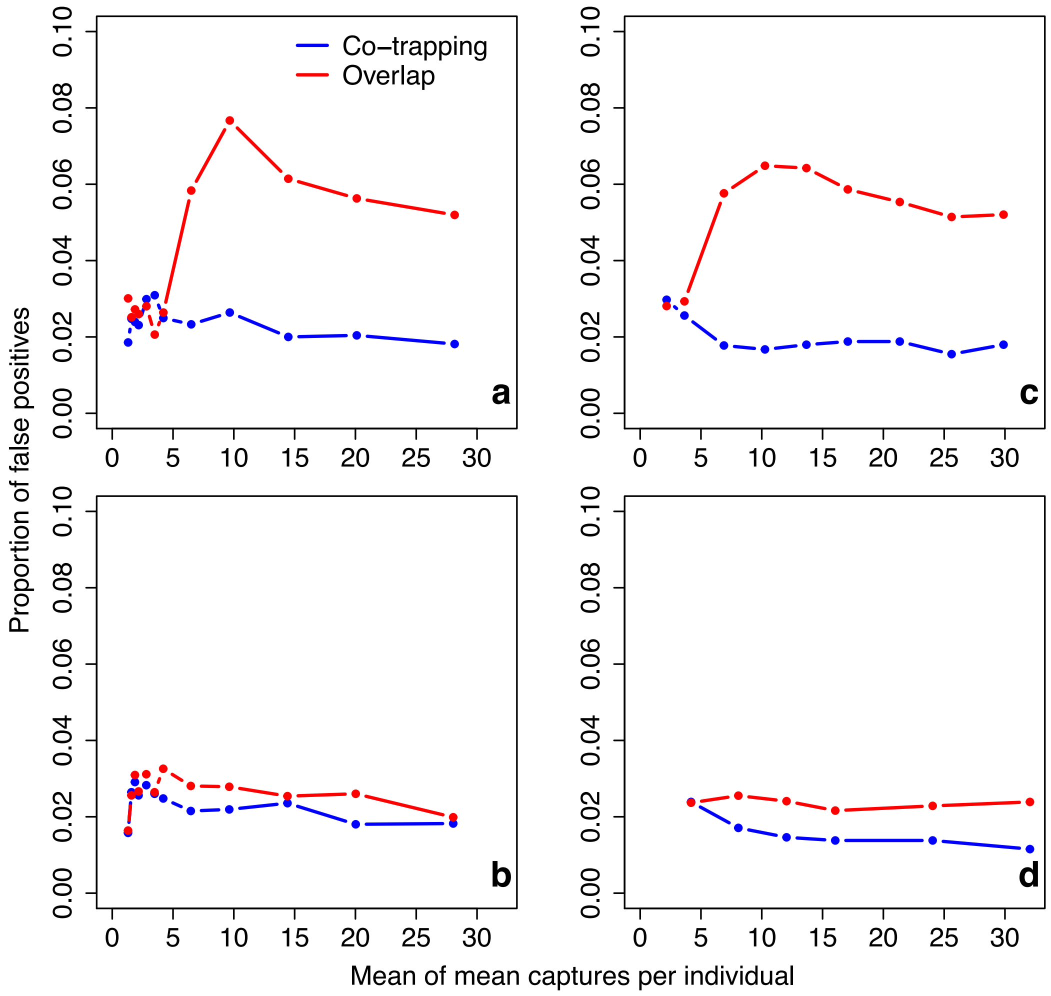


**Fig. S3** Performance of observed networks with varying number of captures per individual on a 10 × 10 trapping grid, as measured by the ability of observed networks to detect a biological effect present when it is not present in the true network. Proportion of false positives shown on *y*-axis, and mean of mean captures per individual shown on *x*-axis. Repeated for (a) observed networks and true networks both generated using our own implementation, (b) observed networks generated using minimum convex polygons (MCPs) and true networks generated using our own implementation, (c) observed networks generated using our own implementation and true networks generated using uniform home ranges, (d) observed networks generated using MCPs and true networks generated using uniform home ranges.

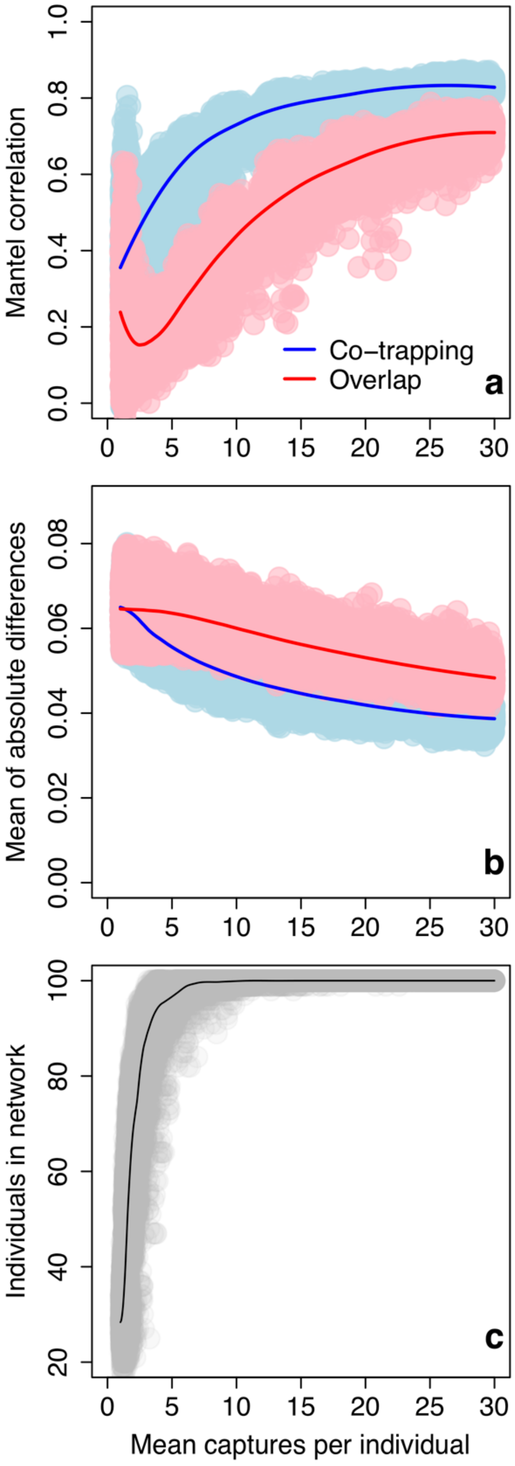


**Fig. S4** Performance of observed networks generated using minimum convex polygons (MCPs) with varying number of captures per individual on a 10 × 10 trapping grid, as measured by (a) Correlation: Mantel correlation between edge weights in observed and true networks generated using our own implementation, (b) Accuracy: Mean of absolute differences in edge weights between observed and true networks generated using our own implementation (lower values = more accurate networks), and (c) Number of individuals in observed networks. LOESS smoother added to aid visual interpretation. Panel (c) refers to the data in the simulated observation dataset, which is identical for both methods.

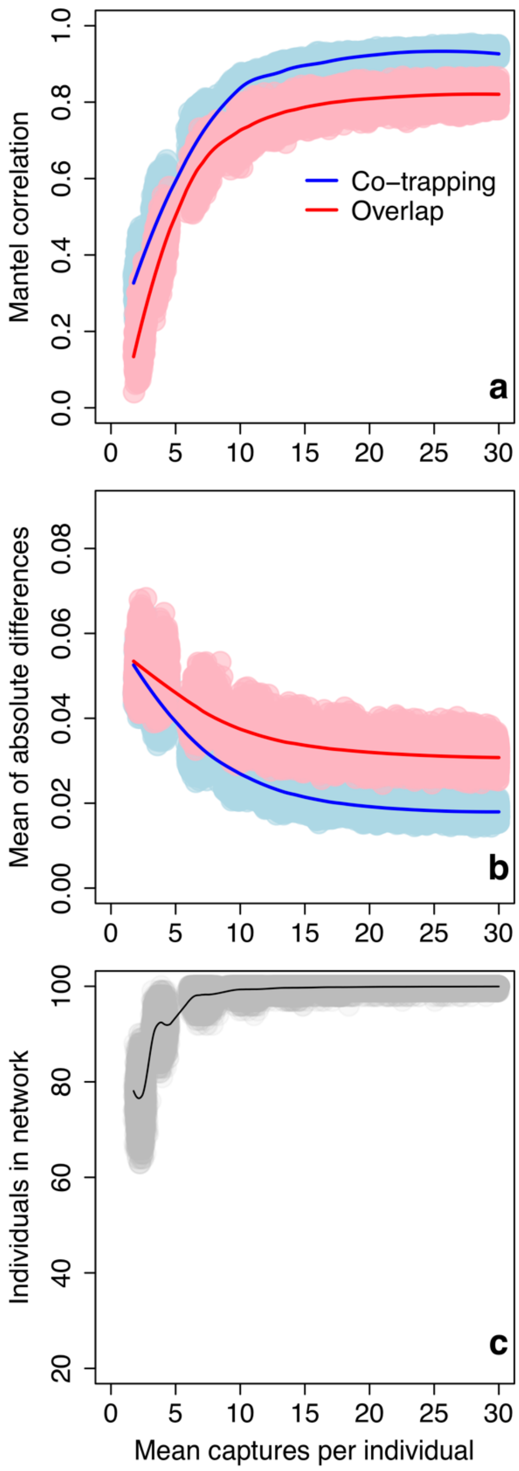


**Fig. S5** Performance of observed networks generated using minimum convex polygons (MCPs) with varying number of captures per individual on a 10 × 10 trapping grid, as measured by (a) Correlation: Mantel correlation between edge weights in observed and true networks generated using uniform home ranges, (b) Accuracy: Mean of absolute differences in edge weights between observed and true networks generated using uniform home ranges (lower values = more accurate networks), and (c) Number of individuals in observed networks. LOESS smoother added to aid visual interpretation. Panel (c) refers to the data in the simulated observation dataset, which is identical for both methods.
